# Predicting continuous ground reaction forces from accelerometers during uphill and downhill running: A recurrent neural network solution

**DOI:** 10.1101/2021.03.17.435901

**Authors:** Ryan S. Alcantara, W. Brent Edwards, Guillaume Y. Millet, Alena M. Grabowski

## Abstract

Ground reaction forces (GRFs) are important for understanding human movement, but their measurement is generally limited to a laboratory. Previous studies used neural networks to predict GRF waveforms during running from wearable device data, but these predictions are limited to the stance phase of level-ground running. We sought to develop a recurrent neural network capable of predicting continuous normal (perpendicular to surface) GRFs across a range of running speeds and slopes from accelerometer data.

19 subjects ran on a force-measuring treadmill at five slopes (0°, ±5°, ±10°) and three speeds (2.5, 3.33, 4.17 m/s) per slope with sacral- and shoe-mounted accelerometers. We then trained a recurrent neural network to predict normal GRF waveforms frame-by-frame. The predicted versus measured GRF waveforms had an average ± SD RMSE of 0.16 ± 0.04 BW and relative RMSE of 6.4 ± 1.5% across all conditions and subjects.

The recurrent neural network predicted continuous normal GRF waveforms across a range of running speeds and slopes with greater accuracy than neural networks implemented in previous studies. This approach may facilitate predictions of biomechanical variables outside the laboratory in near real-time and improves the accurately of quantifying and monitoring external loads experienced by the body when running.

## Introduction

Ground reaction forces (GRFs) are applied to the body when the foot is in contact with the ground and their measurement has facilitated numerous insights into the etiology of running-related injuries^1^. However, measurement of GRFs is generally restricted to a laboratory environment. To determine the effects of sport-specific environments on running kinetics and kinematics, previous studies have replicated aspects of an athlete’s competitive environment (e.g., running surface, slope) within a laboratory environment^2–4^. Alternatively, inertial measurement units (IMUs; wireless wearable devices that measure magnetism, linear acceleration, and angular velocity) have been used to measure athletes’ leg joint angles, stride kinematics, and segmental accelerations during competitive events^5–7^. Although IMUs cannot directly measure GRFs, previous studies have used algorithms to estimate discrete biomechanical variables like peak vertical GRF, ground contact time, vertical impulse, and vertical loading rate^8–12^ from IMU data.

Recently, neural networks have been used to predict GRF waveforms during running^13–16^, from which a variety of discrete variables can be calculated. Although predictions of the entire GRF waveform represent a more versatile outcome compared to predicting a discrete variable, previous studies have used neural network architectures that required waveforms to be normalized to the duration of a step^13^ or stance phase^14,16^, preventing the calculation of biomechanical variables with a temporal component (e.g., ground contact time, step frequency, vertical impulse, and loading rate). Additionally, previous studies have predicted GRF waveforms only during level-ground running^13–16^, limiting the application to environments that are flat (e.g., level treadmill or athletics track). Road and trail running are internationally popular forms of physical activity^17,18^ and require runners to navigate a variety of uphill and downhill slopes. A method that accurately predicts GRF waveforms from wearable device data across a range of running slopes, while maintaining the temporal component, could allow researchers, clinicians, and coaches to measure and monitor a variety of kinetic and kinematic variables in outdoor environments.

Long Short-Term Memory (LSTM) networks^19^ are a type of recurrent neural network that can overcome the traditional requirement of normalizing GRF waveforms to the duration of stance phase because LSTM networks can recurrently predict smaller, uniform portions of a larger sequence of any length. As such, a sequence of continuous GRF data can be predicted if it can be broken up into uniform portions. For the prediction of a given portion, LSTM networks use information from previous portions, effectively “remembering” the portion’s context and have been used to predict sequential data during natural language processing tasks^20^. In the field of Biomechanics, LSTM networks have been used to make frame-by-frame predictions of GRF waveforms using motion capture data^21^ and predictions of the center of mass position relative to center of pressure from IMU data during walking^22^. Developing an LSTM network to predict GRF waveforms from wearable device data would allow researchers to predict GRF waveforms not only during the stance phase, but continuously for the entire duration of a run. IMUs have already been used to longitudinally measure biomechanical variables related to running injury^5–7,10^ and applying an LSTM network to such data could effectively provide a way to indirectly measure continuous GRF waveforms outside of the laboratory at a scale that was previously unattainable.

The purpose of this exploratory study was to develop an LSTM network that could predict continuous normal (perpendicular to running surface) GRF waveforms across a range of running speeds and slopes using data from accelerometers. We sought to develop a network that could predict GRF waveforms with accuracy better than state-of-the-art predictions of time-normalized vertical GRF data during level-ground running using data from multiple IMUs: a root mean square error (RMSE) of 0.21 BW^13^ and relative RMSE (rRMSE; RMSE normalized to the average range of the compared waveforms; Eq. 1) of 13.92%^14^.

## Methods

### Subjects

We analyzed a pre-existing dataset^23–25^ where 21 subjects ran at a combination of running speeds and slopes. Two subjects were excluded from the current analysis due to equipment data acquisition errors, leaving 19 subjects remaining (10 Male, 9 Female; 29 ± 9 years, 173 ± 9 cm, 68.1 ± 9.9 kg). All subjects provided informed consent and the experimental protocol was approved by the University of Calgary Conjoint Health Research Ethics Board (#REB14-1117).

### Experimental Protocol

Following a 5 min warm up at a self-selected speed, each subject completed thirty 30 s trials on a force-measuring treadmill (2000 Hz; Bertec, OH, USA), which included five slopes (0°, ±5°, ±10°) at three speeds (2.5, 3.33, 4.17 m/s) per slope, and three step frequencies (preferred and ± 10%) at 3.33 m/s for each slope. Three custom biaxial accelerometers (2000 Hz) were adhered with tape to subjects during all conditions: two on the right shoe and one on the sacrum. The accelerometers on the shoe were only used to determine the foot strike pattern for each condition using a previously validated method^26^, which provided the percentage of a trial’s foot strikes classified as either rearfoot, midfoot, or forefoot strikes. The biaxial accelerometer on the sacrum was oriented such that it measured vertical and anteroposterior accelerations relative to the accelerometer’s local coordinate system (Figure 1).

**Figure 1 –.**
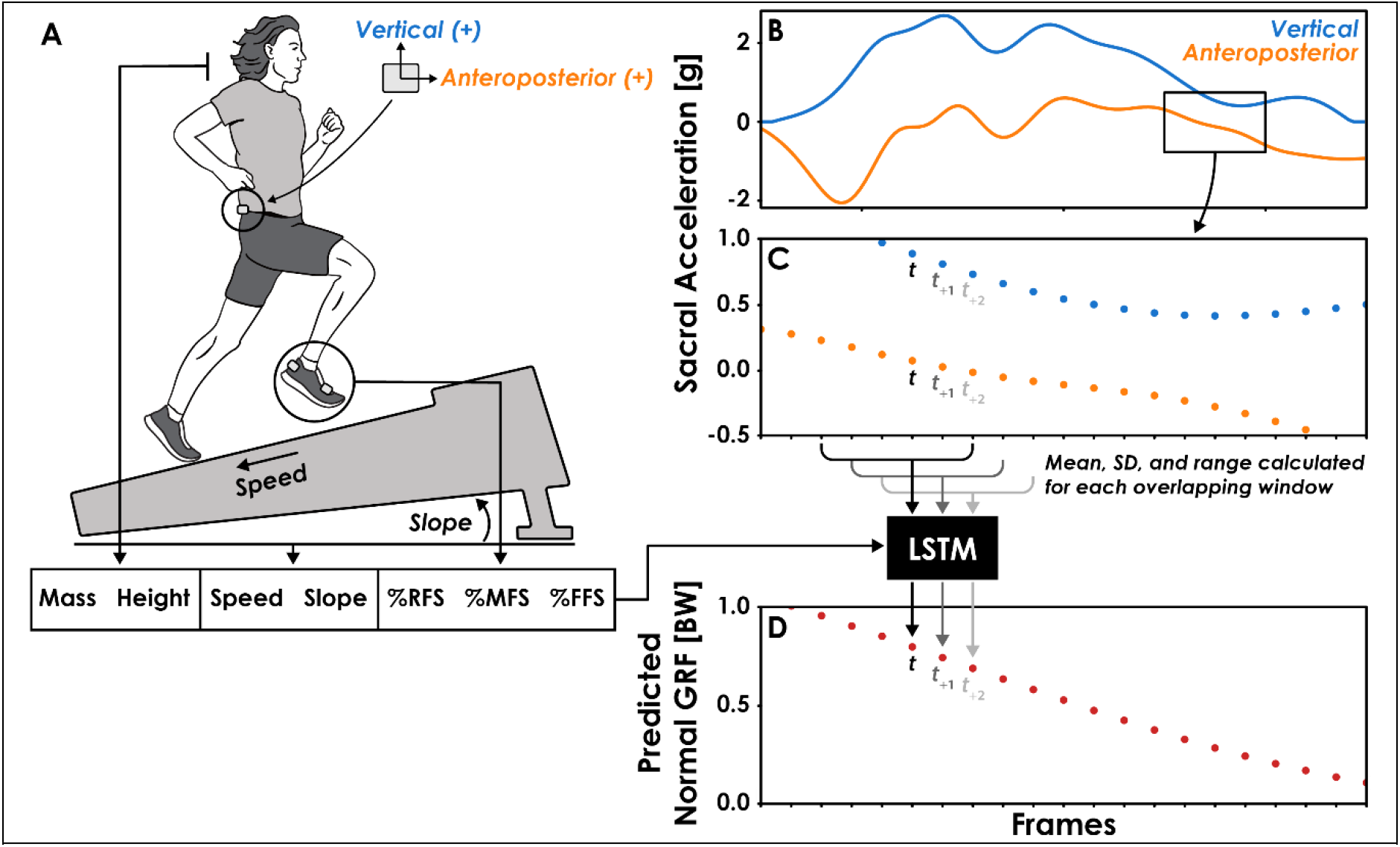
Overview of the Long Short-Term Memory (LSTM) network’s input features and function. **A)** The LSTM network’s input features included body mass, height, running speed, slope, and percentage of a trial’s steps classified as rearfoot (RFS), midfoot (MFS), or forefoot (FFS) strikes. **B)** Filtered vertical and anteroposterior sacral acceleration data, relative to the device’s local coordinate system. **C)** Accelerometer data were divided into overlapping 6 frame (12 ms) windows, one for each frame of the ground reaction force (GRF) data. The mean, standard deviation (SD), and range of vertical and anteroposterior sacral acceleration values were calculated for each window and used as input features to the LSTM network. For the prediction of a normal GRF value at a given time (*t*), the respective window of acceleration data begins at *t*_-3_ and ends at *t*_+2_. **D)** Normal GRFs were predicted frame-by-frame by the LSTM network using the 13 input features.

### Data Processing

We analyzed 5 seconds of data (approximately 13 foot-ground contacts) from each trial and downsampled the normal GRF, vertical sacral acceleration, and anteroposterior sacral acceleration to 500 Hz to reduce the computational cost and match the sampling frequency of prior studies^12,27^. We normalized GRFs to bodyweight (BW) and filtered them using a 4^th^ order low-pass Butterworth filter with a 30 Hz cut-off. We filtered the sacral acceleration data with a 4^th^ order low-pass Butterworth filter with a 20 Hz cut-off. Preliminary analysis revealed that a 20 Hz cut-off improved prediction accuracy and preserved approximately 89% and 82% of the vertical and anteroposterior signal power, respectively.

Vertical sacral accelerometer data were further processed so that all negative values were replaced with zeros. Preliminary analysis revealed that negative vertical acceleration values occur during the aerial phase and replacing them with zeros helped the LSTM network avoid predictions of negative normal GRFs during the aerial phase. For each condition, we used the 2500 frame (5 s trial @ 500 Hz) sequences of vertical and anteroposterior sacral accelerometer data to predict the simultaneously collected 2500 frame sequence of normal GRFs. The recurrent nature of the LSTM network requires sequential data to be divided into smaller portions that are iteratively used to make predictions. To accomplish this, we divided acceleration data for each trial into overlapping windows with a 6 frame (12 ms) width and padded the beginning and end of each trial’s acceleration data with the first and final values, respectively, to ensure the number of windows equaled the number of normal GRF frames (2500) and that the windows were centered on the corresponding frame of the normal GRFs (Figure 1). Pilot testing revealed that a window width of 6 frames was the smallest window we could use without decreasing LSTM network prediction accuracy. Thus, the LSTM network iteratively predicted a single frame of the normal GRF at time *t* using acceleration data from frames *t*_-3_ through *t*_+2_ (Figure 1).

### Feature Engineering

A total of 13 features were used as inputs in the LSTM network (Figure 1). We calculated the mean, standard deviation (SD), and range of vertical and anteroposterior acceleration data for each 12 ms window and used them as input features. The use of summary statistics as input features has been shown to maintain neural network accuracy while benefiting from a reduced computational cost^28^. These three summary statistics were normalized to a range of 0 – 1 and represent 6 (3 features x 2 acceleration axes) of the 13 input features. The remaining input features were selected due to their effect on running kinetics and kinematics: subject height, body mass, running speed, slope, and percentage of steps classified as either rearfoot, midfoot, or forefoot strikes^12,24,25,29,30^. We chose not to include step frequency as an input feature, despite the presence of the ±10% preferred step frequency conditions, to increase the variability in the data used to predict GRF waveforms. Doing so theoretically represents a greater challenge for the LSTM network as there is additional variability between trials that is not being explicitly accounted for with an input variable.

### Neural Network Architecture

The neural network consisted of a Bidirectional LSTM and a multilayer perceptron (MLP) with three fully connected layers containing 128, 384, and 320 neurons, respectively (Figure 2). The Bidirectional LSTM consists of two LSTM layers where the order of the input sequence is reversed for the second layer. Reversing the sequence for the second LSTM layer allows the network to utilize information from future portions of the sequence just as the first LSTM layer utilizes information from prior portions. The outputs from each LSTM layer are then averaged before being passed along to the MLP. The number and size of the layers were determined using the Hyperband hyperparameter optimization algorithm^31^ on the data of two randomly selected subjects. The LSTM network was trained using a batch size of 32, learning rate of 0.001, and mean square error loss function. Network weights and biases were updated using the adaptive moment estimation (Adam) optimization algorithm at the end of each epoch^32^ and training lasted a maximum of 1000 epochs or until the mean square error failed to decrease by 0.001 BW after 30 consecutive epochs. The neural network was developed using the Tensorflow (v2.2.0) python library^33^.

**Figure 2 –.**
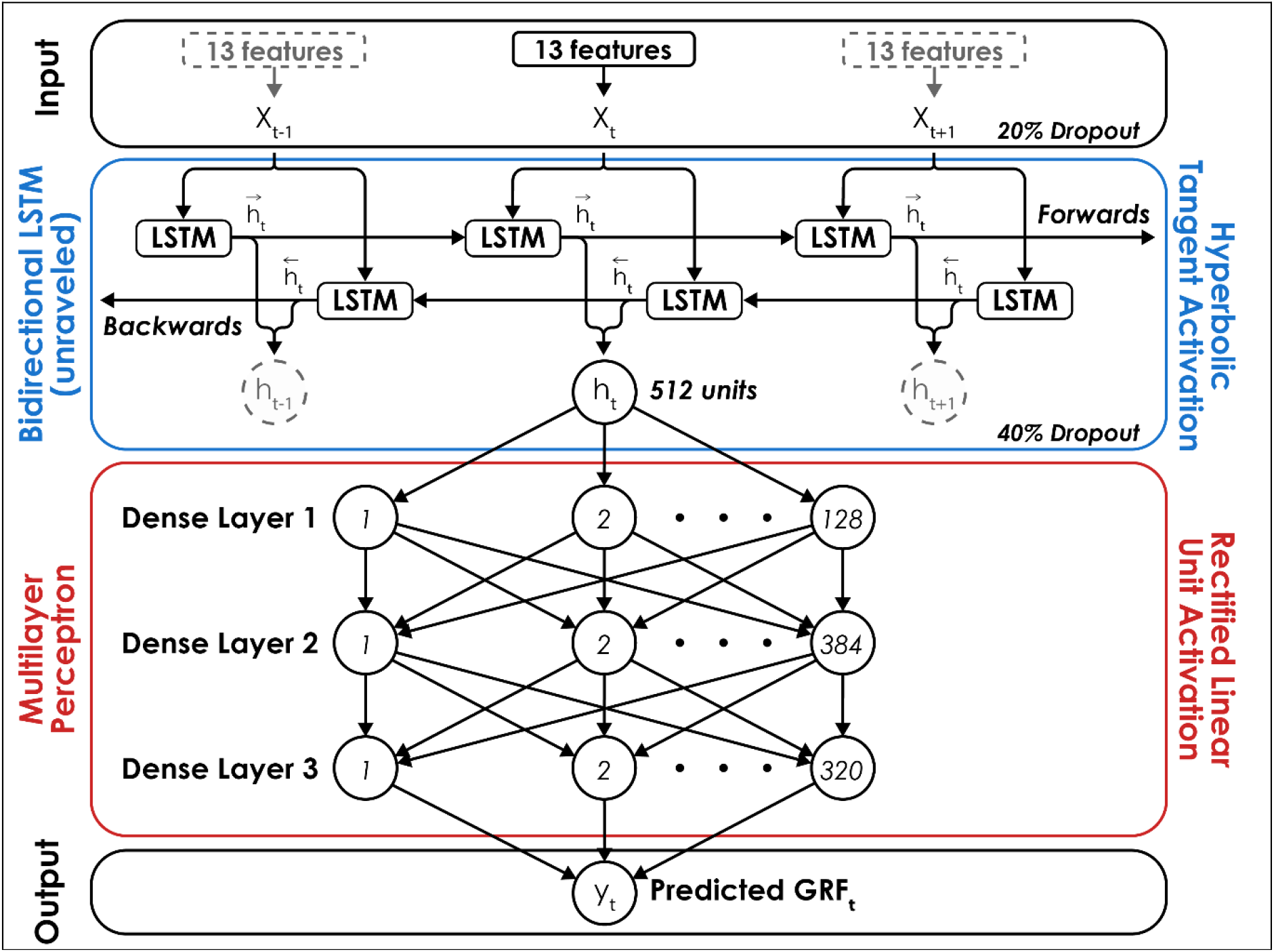
The Long Short-Term Memory (LSTM) network consisted of a Bidirectional LSTM layer with a hyperbolic tangent activation function followed by a multilayer perceptron (MLP) with rectified linear unit activation functions for three hidden layers with 128, 384, and 320 neurons, respectively. The Bidirectional LSTM layer is unraveled to illustrate its recurrent nature and dashed lines signify inputs (*x*) and outputs (*h*) at time *t*_-1_ and *t*_+1_. A dropout rate of 20% was applied to the input layer of the network and a dropout rate of 40% was applied to the output of the Bidirectional LSTM layer to limit network overfitting. For each prediction of the normal ground reaction force (GRF) at a given time (*t*), the network received 13 features as inputs (*x*_*t*_; Figure 1), passed the output from the Bidirectional LSTM layer (*h*_*t*_) to the MLP, and predicted a single value (*y*_*t*_) with a linear activation function in the output layer.

### Network Validation

We assessed the accuracy and generalizability of the network using a Leave-One-Subject-Out (LOSO) cross validation method^34^. LOSO cross validation is a variation of K-fold cross validation that requires the dataset to be subset by subject, with one subject’s data withheld for testing purposes and the rest of the subjects’ data used to train the network. This process is repeated until the network has been tested on every subject’s data, ultimately providing an ensemble of networks and their respective accuracy metrics. Performing LOSO cross validation can be computationally costly, as the network must be trained and tested a number of times equal to the number of subjects (n = 19), but the benefits of this method include the ensemble of accuracy metrics and assurance that a given subject’s data are not included in both the training and testing subsets, which can artificially increase the reported accuracy of a network^35,36^.

In addition to the LOSO cross validation method, we performed a test-train split of one representative subject’s data according to slope (±5° trials reserved for testing, 0° and ±10° slopes used for training) to test the accuracy of the model when predicting speed-slope combinations that were not present during training. We selected Subject 14 as a representative subject because their RMSE during LOSO cross validation was similar to the average RMSE across all subjects (Figure 3) and their GRF waveforms illustrated an interaction between running slope and normal GRF impact peak magnitude^37^. This single-subject validation method prioritizes accuracy over generalizability (ability to make accurate predictions for a variety of individuals) and represents a potential circumstance where an LSTM network is trained on data collected from a single athlete prior to their competitive season and later used to predict only that athlete’s GRF data from wearable device data during their competitive season.

**Figure 3 –.**
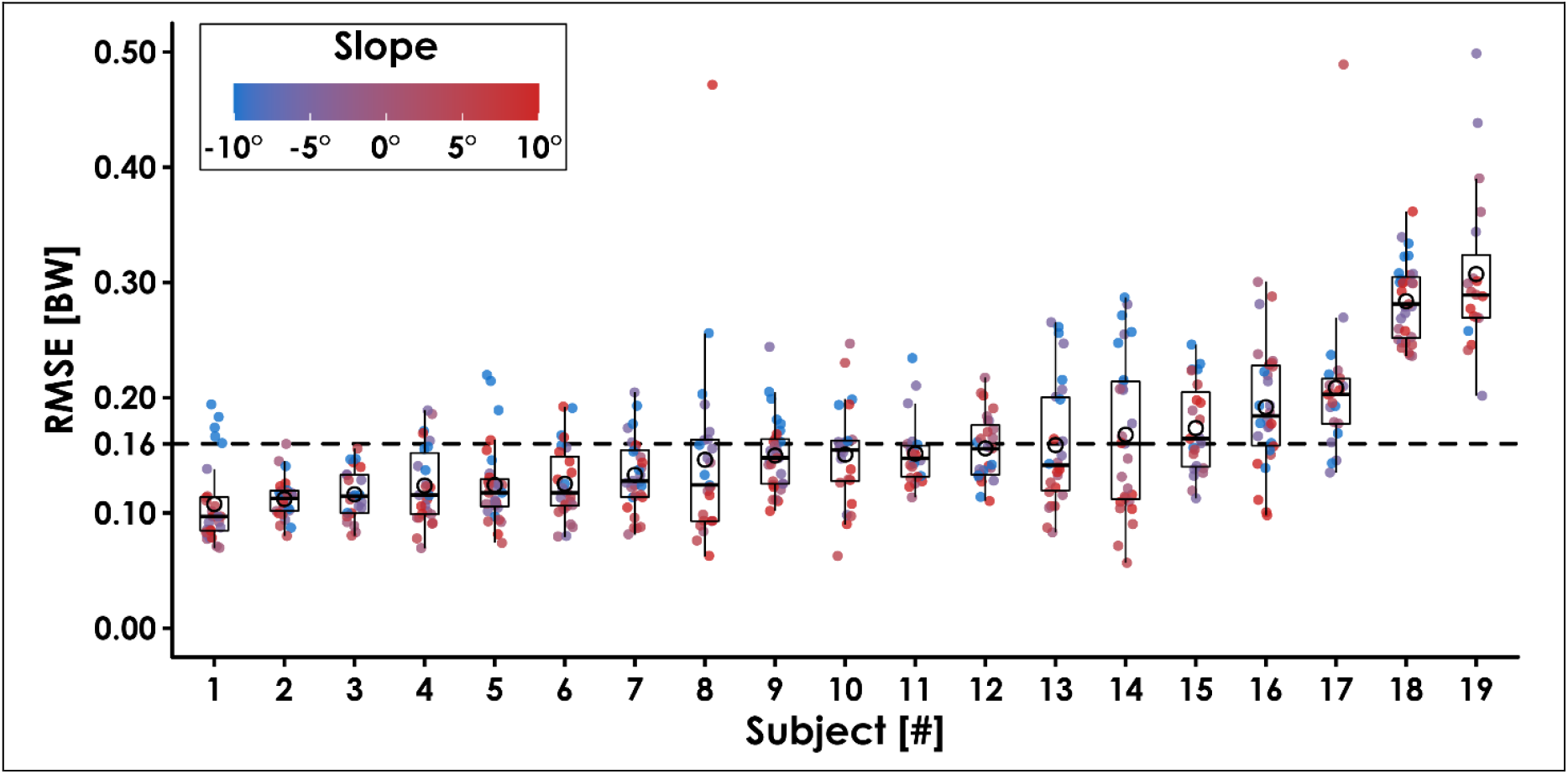
Ground reaction force waveform prediction error for each subject across all conditions. The average root mean square error (RMSE) across subjects was 0.16 BW (dotted line). Filled circles represent each trial, and the color indicates slope (0°, ±5°, ±10°) at three speeds (2.5, 3.33, 4.17 m/s). Open circles represent each subject’s average RMSE, horizontal bars are the median RMSE, box plot edges indicate the interquartile range (IQR; 25^th^ and 75^th^ percentile), and the whiskers encompass values that fall within 1.5*IQR. Subjects are sorted from lowest to highest RMSE.

Prediction error for each trial’s GRF waveforms was quantified as the root mean square error (RMSE) and relative RMSE (rRMSE), which is RMSE normalized to the average range of compared waveforms, expressed as a percentage and defined as

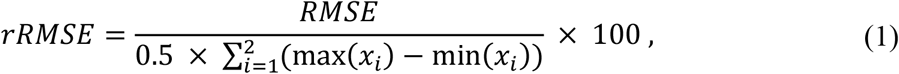

where *x*_1_ and *x*_2_ are the GRF waveforms predicted by the LSTM network and measured by the force-measuring treadmill^38^. Additionally, we used a threshold of 5% BW to identify stance phase and calculated the active peak of the normal GRF waveform, normal impulse, normal GRF loading rate, contact time, and step frequency from the predicted and measured GRF data. The normal GRF active peak was calculated as the maximum normal GRF value occurring between 40 – 60% of stance phase because the magnitude of the impact peak can exceed the active peak during downhill running and occurs during early stance phase (0 – 30%)^25,37^. We calculated impulse as the integral of the normal GRF waveform during the stance phase with respect to time, loading rate as the average slope of the normal GRF waveform during the first 25 ms of stance phase^39^, contact time as the duration when the normal GRF was greater than 5% BW, and step frequency as the number of initial foot-ground contacts per second. We report the mean absolute percent error (MAPE) of these discrete variables for each subject. Data analysis was performed in python (v3.6.9) and R (v4.0.4) using custom libraries^40–47^.

We enforced two biomechanical boundaries upon the predicted GRF waveforms to ensure that data fell within established biomechanical limits and could be used to calculate the discrete biomechanical variables of interest. First, the predicted GRF waveform had to have an equal number of foot-ground contacts as the GRF waveform measured by the force-measuring treadmill, determined using the same 5% BW threshold. Second, the step frequency over the duration of the predicted GRF waveform had to be ≤ 4 Hz. We selected these criteria based on previous research of running biomechanics, as thresholds of 5% BW have been previously used to identify the stance phase for the calculation of kinetic or kinematic variables^12,27^ and during uphill and downhill running, step frequency is ≤ 4 Hz^48,49^. Trials that failed to meet either of these criteria were used to calculate the LSTM network’s overall prediction failure rate and removed from subsequent analyses.

## Results

We analyzed 529 trials for the present study. The predicted GRF waveforms for 32 trials (6%) failed to meet one or both criteria and were considered failed predictions. Specifically, we identified 22 trials (4%) that required a threshold greater than 5% BW to identify an equal number of steps between predicted and measured GRF waveforms and 10 trials (2%) that had a step frequency greater than 4 Hz. Thus, 94% of GRF waveforms predicted by the LSTM network fell within the imposed biomechanical boundaries.

Leave-One-Subject-Out cross validation revealed that the LSTM network predictions of each subject’s normal GRF waveforms had an average ± SD RMSE of 0.16 ± 0.04 BW (Figure 3) and rRMSE of 6.4 ± 1.5% compared to GRF waveforms measured by the force-measuring treadmill across all conditions (Table 1). RMSE values were generally lower during slow uphill running (2.5 m/s, +10°; 0.13 BW) compared to fast downhill running (4.17 m/s, -10°; 0.20 BW) (Figures 4 and 5). The MAPE for step frequency was 0.1 ± 0.1%, contact time was 4.9 ± 4.0%, impulse was 6.4 ± 6.9%, normal GRF active peak was 8.5 ± 8.2%, and loading rate was 27.6 ± 36.1% (Table 2).

**Table 1.** Mean ± SD root mean square error (RMSE) and relative RMSE (rRMSE) for normal GRF waveforms predicted by the LSTM network compared to the measured normal GRF waveforms for each subject.

| Subject | RMSE [BW] | rRMSE [%] |
| --- | --- | --- |
| 1 | 0.11 ± 0.04 | 4.0 ± 1.1 |
| 2 | 0.11 ± 0.02 | 4.1 ± 0.7 |
| 3 | 0.12 ± 0.02 | 4.4 ± 0.9 |
| 4 | 0.12 ± 0.03 | 4.6 ± 1.2 |
| 5 | 0.12 ± 0.04 | 4.7 ± 0.9 |
| 6 | 0.12 ± 0.03 | 5.2 ± 1.4 |
| 7 | 0.13 ± 0.03 | 5.1 ± 1.2 |
| 8 | 0.15 ± 0.08 | 5.3 ± 3.3 |
| 9 | 0.15 ± 0.03 | 5.4 ± 1.0 |
| 10 | 0.15 ± 0.04 | 5.2 ± 1.3 |
| 11 | 0.15 ± 0.03 | 5.8 ± 0.8 |
| 12 | 0.16 ± 0.03 | 6.1 ± 1.1 |
| 13 | 0.16 ± 0.06 | 5.7 ± 1.5 |
| 14 | 0.17 ± 0.07 | 6.9 ± 2.5 |
| 15 | 0.17 ± 0.04 | 7.9 ± 2.0 |
| 16 | 0.19 ± 0.05 | 6.7 ± 1.7 |
| 17 | 0.21 ± 0.07 | 7.2 ± 2.5 |
| 18 | 0.29 ± 0.03 | 14.0 ± 1.7 |
| 19 | 0.31 ± 0.07 | 13.7 ± 2.7 |
| Mean ± SD | 0.16 ± 0.04 | 6.4 ± 1.5% |

**Table 2.** Mean ± SD discrete variables calculated from normal ground reaction force (GRF) waveforms predicted by the LSTM network (“Predicted”) and normal GRF waveforms measured from the force-measuring treadmill (“Measured”) across all speeds and subjects for each slope.

| **Slope** | **Step Frequency [Hz]** |  | **Contact Time [ms]** |  | **Impulse [BW\*s]** |  |
| --- | --- | --- | --- | --- | --- | --- |
|  | Predicted | Measured | Predicted | Measured | Predicted | Measured |
| -10° | 3.1 ± 0.3 | 3.1 ± 0.3 | 215 ± 26 | 214 ± 29 | 0.33 ± 0.03 | 0.33 ± 0.05 |
| -5° | 3.1 ± 0.3 | 3.1 ± 0.3 | 222 ± 23 | 223 ± 27 | 0.34 ± 0.04 | 0.34 ± 0.04 |
| 0° | 3.1 ± 0.3 | 3.1 ± 0.3 | 228 ± 24 | 229 ± 27 | 0.33 ± 0.04 | 0.34 ± 0.04 |
| +5° | 3.2 ± 0.3 | 3.2 ± 0.3 | 228 ± 25 | 232 ± 26 | 0.32 ± 0.03 | 0.33 ± 0.04 |
| +10° | 3.4 ± 0.3 | 3.4 ± 0.3 | 223 ± 26 | 227 ± 26 | 0.30 ± 0.04 | 0.30 ± 0.04 |

| **Slope** | **Active Peak [BW]** |  | **Loading Rate [BW/s]** |  |
| --- | --- | --- | --- | --- |
|  | Predicted | Measured | Predicted | Measured |
| -10° | 2.38 ± 0.24 | 2.36 ± 0.40 | 64.7 ± 19.9 | 68.4 ± 20.6 |
| -5° | 2.41 ± 0.25 | 2.45 ± 0.36 | 54.9 ± 16.7 | 60.6 ± 19.4 |
| 0° | 2.42 ± 0.22 | 2.51 ± 0.36 | 42.0 ± 12.8 | 49.0 ± 17.2 |
| +5° | 2.36 ± 0.21 | 2.40 ± 0.33 | 34.8 ± 8.5 | 39.3 ± 13.3 |
| +10° | 2.24 ± 0.22 | 2.20 ± 0.29 | 30.1 ± 8.3 | 31.3 ± 12.7 |

**Figure 4 –.**
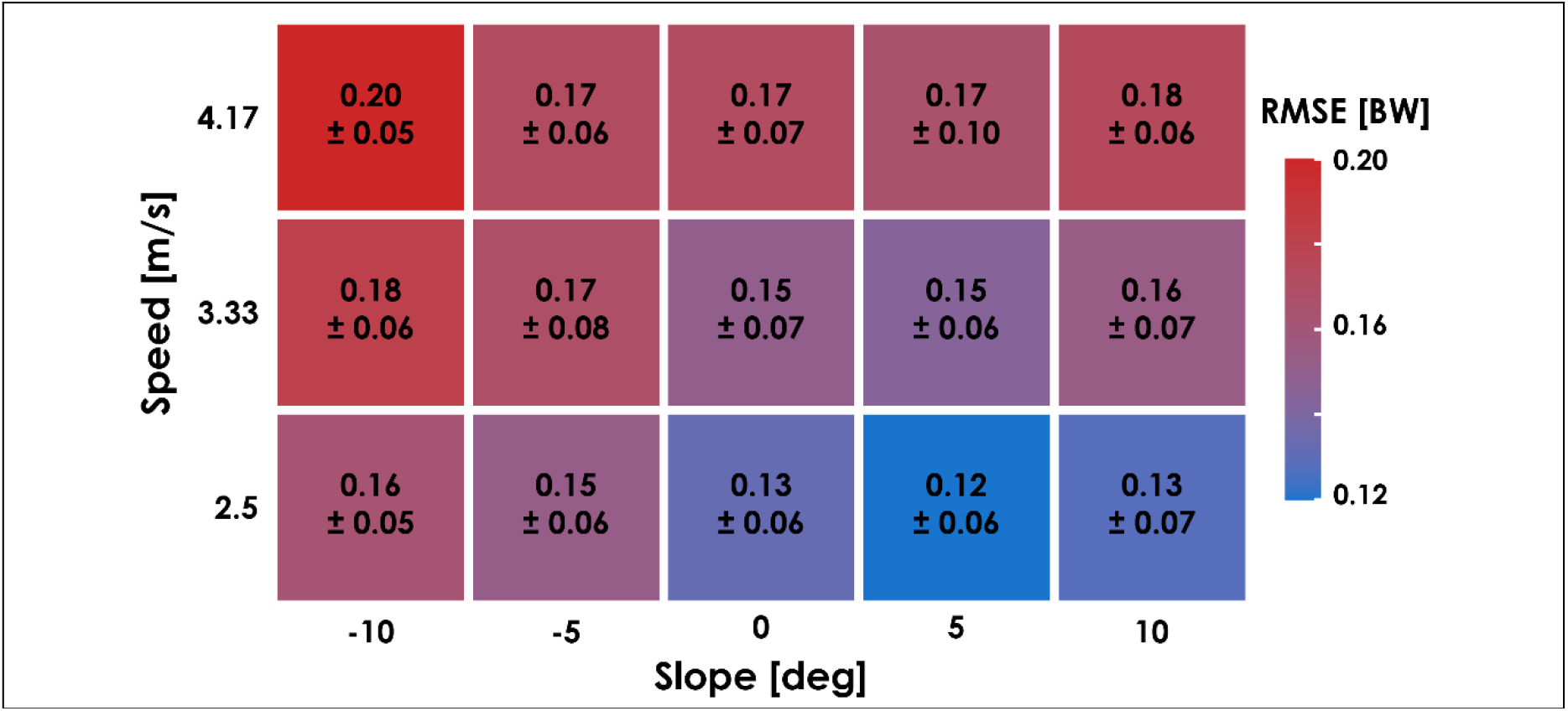
The average ± SD root mean square error (RMSE) of the predicted ground reaction force (GRF) waveforms compared to the GRF waveform measured by the force-measuring treadmill for each condition during leave-one-subject-out (LOSO) cross validation.

**Figure 5 –.**
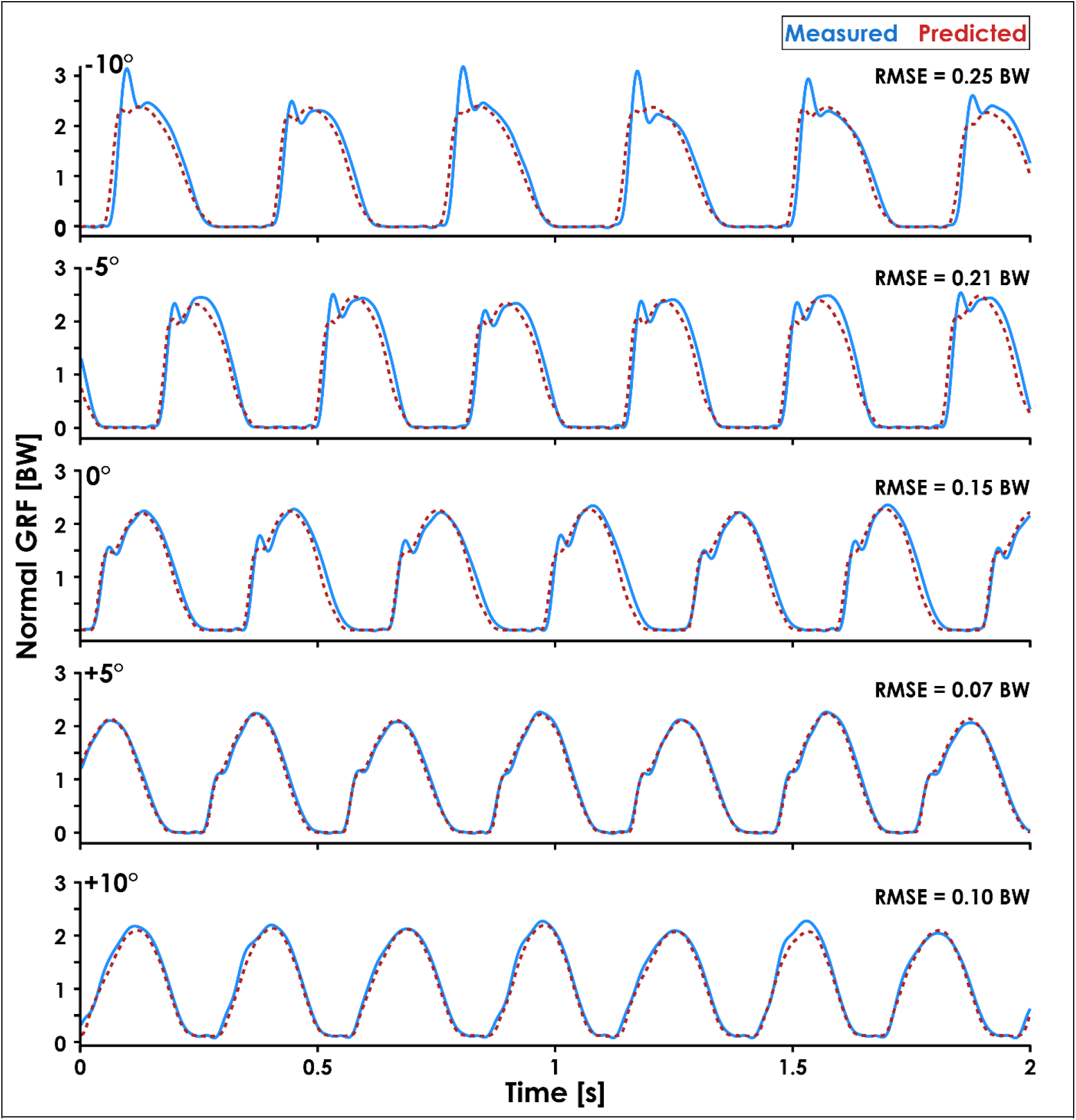
Normal ground reaction force (GRF) waveforms predicted by the recurrent neural network (dashed red lines) and measured by the force-measuring treadmill (solid blue lines) at 3.33 m/s and all slopes (0°, ±5°, ±10°) are presented for Subject 14. Subject 14 was selected because they had similar RMSE values (0.17 ± 0.07 BW) as the average across all subjects (0.16 ± 0.04 BW) and their GRF waveforms illustrate an interaction between running slope and normal GRF impact peak magnitude.

The prediction error for one representative subject’s (Subject 14) normal GRF waveforms at ±5° during single-subject validation was lower than those resulting from the LOSO cross validation, with an average ± SD RMSE of 0.08 ± 0.02 BW and rRMSE of 3.3 ± 0.9%. The MAPE of step frequency (0.1 ± 0.1 %), contact time (3.0 ± 2.3%), impulse (2.5 ± 1.9%), normal GRF active peak (2.7 ± 2.0%), and loading rate (17.6 ± 16.9%) calculated from predicted GRF waveforms were also generally lower than those resulting from LOSO cross validation.

## Discussion

We developed a recurrent neural network capable of predicting continuous normal GRF waveforms across a range of running speeds (2.5 – 4.17 m/s), slopes (0°, ±5°, ±10°), and step frequencies (preferred, ±10%) from accelerometer data. Our findings indicate that an LSTM network given the runner’s mass, height, running speed, slope, foot strike pattern, and sacral acceleration can predict normal GRF waveforms across a range of speeds and slopes with an RMSE of 0.12 – 0.20 BW and rRMSE of 5.4 – 7.3% (Figure 4). For comparison, previous studies report an RMSE of 0.39 ± 0.26 BW^16^, an RMSE of 0.21 ± 0.03 BW^13^, and an rRMSE of 13.92%^14^ when using neural networks to predict the stance phase vertical GRF waveform during level-ground running. In contrast to previous studies, the LSTM network does not require preliminary stance phase identification or time normalization, which preserves the temporal component of the predicted GRF waveform. This characteristic of the LSTM network allowed us to calculate stride kinematic variables like step frequency and contact time with a MAPE < 5%. Additionally, the recurrent nature of the LSTM network facilitates frame-by-frame predictions of GRF waveforms and can be used to make predictions over any duration of running. Thus, an LSTM network could be used to quantify changes in normal GRF waveforms over the course of a prolonged run (e.g., a marathon race).

The accuracy of predicted GRF waveforms varied across speeds and slopes, with a combination of faster running speeds and negative slopes producing greater RMSE values than slower running speeds and positive slopes (Figure 4). The greater RMSE values during downhill running may be due to the LSTM network’s inability to account for changes in impact peak magnitude across slopes (Figure 5). Previous studies have found that the presence of an impact peak in the normal GRF waveform is subject-specific, affected by changes in running slope, and associated with acceleration of the effective mass of the lower extremity during early stance phase^25,37,50^. Thus, predictions of normal GRF waveforms across slopes may be further improved by incorporating accelerations measured at the feet or lower legs.

We also quantified the accuracy of the LSTM network when trained and tested on data from the same subject. Although not a valid method of determining the LSTM network’s generalizability, single-subject validation provides insight into how well a personalized neural network could predict an individual’s GRF waveforms for unknown combinations of speed and slope in the future. We found that predicted GRF waveforms of a representative subject (Subject 14) during the ±5° slope conditions had an average ± SD RMSE of 0.08 ± 0.02 BW, compared to 0.16 ± 0.03 BW during LOSO cross validation. These findings indicate that a subject-specific LSTM network was twice as accurate as the LOSO cross validated LSTM network. A single-subject approach may be particularly beneficial for researchers, coaches, or clinicians who have the resources to train personalized LSTM networks and wish to monitor a specific athlete’s biomechanics over the course of a competitive season. For example, an athlete could run at a variety of speeds and slopes while wearing accelerometers during a baseline data collection on a force-measuring treadmill at the start of their competitive season and a personalized LSTM network could be trained on their data. Then, if accelerometer data were collected from an athlete during training runs, their normal GRF waveforms and a variety of discrete values could be predicted and monitored longitudinally.

The MAPE values for step frequency, contact time, impulse, and normal GRF active peak were ≤ 8.5%, but the loading rate MAPE was 27.6 ± 36.1%. The lower MAPE values for step frequency, contact time, impulse, and normal GRF active peak indicate that the LSTM network consistently identified the general shape of the GRF waveform and the boundaries of the stance phase despite changes in speed, slope, and step frequency. However, the network did not consistently predict the presence of an impact peak during early stance phase (Figure 5, -10° trial), which affected the predicted slope of the GRF waveform during early stance phase and thus the accuracy of loading rate values. Although the prominence of an impact peak in the normal GRF waveform is affected by foot strike pattern and slope^37^, two of the inputs for the LSTM network, the decreased accuracy when estimating loading rate may be because we did not include accelerometer data from the lower legs or feet. A previous study found moderate-strong correlations between axial tibial acceleration and vertical GRF impact peak magnitude (r = 0.76) and timing (r = 0.94) during running^51^. We did not include accelerometer data from the shoes as inputs for the LSTM network because the data were not available for both feet.

Recurrent neural networks represent a promising strategy for predicting continuous normal GRFs from wearable devices in outdoor environments. The LSTM network required data from three accelerometers (one on the sacrum and two on the right shoe to determine foot strike pattern), but we also performed a *post-hoc* analysis of prediction accuracy without the foot strike pattern data to quantify the network’s accuracy when only using data from one sacral accelerometer. The *post-hoc* analysis revealed that excluding foot strike pattern data slightly increased the average ± SD RMSE from 0.16 ± 0.04 BW to 0.17 ± 0.05 BW and rRMSE from 6.4 ± 1.5% to 6.7 ± 1.7%. Excluding foot strike pattern data affected the MAPE of discrete variables by < 3% (Table 3). These findings indicate that the LSTM network can predict normal GRF waveforms from a single accelerometer on the sacrum more accurately than neural networks implemented in previous studies (RMSE = 0.21 – 0.39 BW, rRMSE = 13.92%), which required data from 3 – 7 wearable devices^13,14,16^.

**Table 3.**
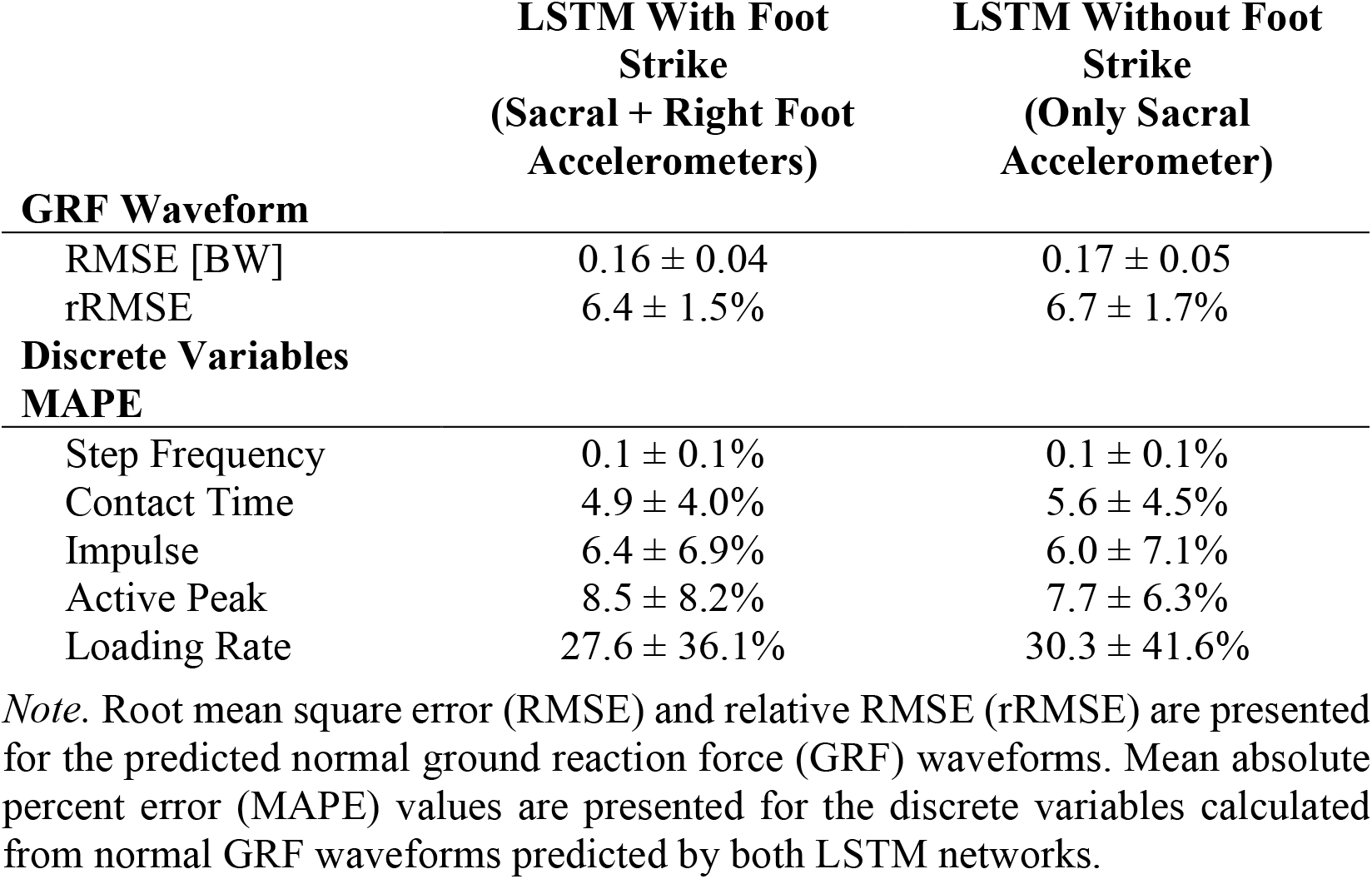
Error metrics for the predicted waveforms and discrete variables when training the Long Short-Term Memory (LSTM) network with and without foot strike pattern as an input feature.

Using a recurrent neural network in combination with accelerometers and a global positioning system (GPS) device to obtain speed and slope data could potentially allow runners to receive biomechanical feedback during an outdoor run. Watches with GPS capabilities are commonly used by runners^52^, have been used to provide real-time feedback of step frequency^53^, and could provide running speed and slope data to the LSTM network to predict GRF waveforms in near-real time^54^. Discrete biomechanical variables could then be calculated from predicted normal GRF waveforms and sent to a clinician, coach, researcher, or the runner themselves. A similar approach has been implemented during outdoor walking and running using an integrated IMU-GPS device placed in a backpack, but it is unclear how accurate or generalizable this approach is as the network was trained and tested on data from three subjects and the reported accuracy metrics were combined for walking and running^55^. To facilitate calculation of GRF-based variables during outdoor running using accelerometers, we have made the LSTM networks, which were trained on all subjects, with and without the need for foot strike data, publicly available alongside a tutorial at www.github.com/alcantarar/Recurrent_GRF_Prediction.

There are potential limitations to consider alongside our findings. The accelerometers used in the present study were adhered to subjects using tape and a less secure attachment method may introduce movement artefact into the accelerometer data. Previous research suggests that attachment method can affect peak tibial acceleration during running^56^, but the lower leg experiences larger accelerations than the sacrum during running^23^ and thus is more sensitive to different attachment methods. Using the LSTM network to predict normal GRF waveforms from a sacral accelerometer adhered differently than in the present study may affect prediction accuracy, but the 20 Hz low-pass filter we applied to the accelerometer data can potentially mitigate this effect. Additionally, predictions made with the LSTM network may not be generalizable for speeds or slopes that fall outside the range of the training data (2.5 – 4.17 m/s and ±10°) as biomechanics change when running on steep slopes (e.g. 20 – 40°)^3,57^ or with changes in speed^12^. Lastly, the LSTM network was trained on data collected on a stiff force-measuring treadmill and thus accelerometer data collected during running on less stiff surfaces (e.g., grass) may result in greater prediction errors given the effects of surface stiffness on running biomechanics and thus energy absorption^58,59^.

In conclusion, we developed a recurrent neural network that used accelerometer data to predict continuous normal GRF waveforms across a range of running speeds (2.5 – 4.17 m/s) and slopes (0°, ±5°, ±10°) with an average ± SD RMSE of 0.16 ± 0.04 BW and rRMSE of 6.4 ± 1.5%. Unlike neural networks implemented in prior studies, the recurrent neural network does not require preliminary identification of the stance phase or temporal normalization and allows for near real-time predictions of normal GRF waveforms during running. Accurate predictions of normal GRF waveforms using wearable devices will improve the ability to longitudinally monitor biomechanical variables in non-laboratory environments.

## Acknowledgements

We utilized resources from the University of Colorado Boulder Research Computing Group, which is supported by the National Science Foundation (awards ACI-1532235 and ACI-1532236), the University of Colorado Boulder, and Colorado State University.

## Notes

**Conflict of Interest Disclosure:** None

https://github.com/alcantarar/Recurrent_GRF_Prediction

### Competing Interest Statement

The authors have declared no competing interest.

### Summary of Updates

Cut down wordcount.

